# Impaired KCC2 phosphorylation leads to neuronal network dysfunction and neurodevelopmental pathogenesis

**DOI:** 10.1101/606566

**Authors:** Lucie I. Pisella, Jean-Luc Gaiarsa, Diabé Diabira, Jinwei Zhang, Ilgam Khalilov, JingJing Duan, Kristopher T. Kahle, Igor Medina

**Affiliations:** Aix-Marseille University UMR 1249, INSERM (Institut National de la Santé et de la Recherche Médicale) Unité 1249, INMED (Institut de Neurobiologie de la Méditerranée), Marseille, France; Institute of Biomedical and Clinical Sciences, College of Medicine and Health, University of Exeter, Hatherly Laboratories, Exeter, EX4 4PS, UK; Department of Neurobiology, Howard Hughes Medical Institute, Boston Children’s Hospital, Boston, Massachusetts 02115, USA; Departments of Neurosurgery, Pediatrics, and Cellular & Molecular Physiology; and Centers for Mendelian Genomics, Yale School of Medicine, New Haven, CT 06510, USA

**Author notes:** Corresponding authors: Dr. Igor Medina (INMED / INSERM), Dr. Kristopher Kahle (Yale University).

## Abstract

KCC2 is a vital neuronal K^+^/Cl^-^ co-transporter that is implicated in the etiology of numerous neurological diseases. It is subject to developmental dephosphorylation at threonine 906 and 1007, the functional importance of which remains unclear. We engineered mice with heterozygous phospho-mimetic mutations T906E and T1007E (*KCC2*^*E/+*^) to prevent the normal developmental dephosphorylation of these sites. Immature (P15) but not juvenile (P30) *KCC2*^*E/+*^ mice exhibited altered GABAergic inhibition, an increased glutamate/GABA synaptic ratio, and higher seizure susceptibility. *KCC2*^*E/+*^ mice also had abnormal ultra-sonic vocalizations at P10-P12 and impaired social behavior at P60. Post-natal bumetanide treatment restored network activity at P15 but not social behavior at P60. Our data show that post-translational KCC2 regulation controls the GABAergic developmental sequence *in vivo*. The post-translational deregulation of KCC2 could be a risk factor for the emergence of neurological pathology and the presence of depolarizing GABA is not essential for manifestation of behavioral changes.

## INTRODUCTION

Neurodevelopmental disorders (NDDs), including specific epilepsy subtypes, schizophrenia, intellectual disability, and autism spectrum disorders (ASDs), exhibit a shared spectrum of pathological changes in neuronal morphology, synapse function, and network properties (Deidda *et al.*, 2014; Pasciuto *et al.*, 2015; Cattane *et al.*, 2018). A fundamental obstacle to developing novel therapies for NDDs is our limited knowledge of disease pathogenesis. Recent data has implicated impaired function of the neuronal potassium/chloride extruder KCC2 in the pathogenesis of multiple NDDs (Schulte *et al.*, 2018).

KCC2 (*SLC12A5*) is essential for establishing and maintaining the low intracellular chloride concentration ([Cl^-^]_i_) of mature mammalian neurons. The link between KCC2 and NDDs has been highlighted by several studies showing a decreased functional expression of KCC2 in human patients with epilepsy (Kahle *et al.*, 2016*b*), schizophrenia (Hyde *et al.*, 2011; Sullivan *et al.*, 2015), and Rett syndrome (Duarte *et al.*, 2013), as well as in mouse models of Fragile X (Tyzio *et al.*, 2014) and Rett syndrome diseases (El-Khoury *et al.*, 2014; Banerjee *et al.*, 2016). However, due to the post-translational regulation of KCC2, there is no simple relationship between the abundance of KCC2 protein and its activity as chloride extruder (Kahle *et al.*, 2013). Phosphorylation of KCC2 at specific residues strongly affects the ion transport activity of the transporter (Lee *et al.*, 2007, 2010; Inoue *et al.*, 2012; Weber *et al.*, 2014; Friedel *et al.*, 2015; Cordshagen *et al.*, 2018).

Among multiple KCC2 phosphorylation sites (Payne *et al.*, 1996; Lee *et al.*, 2007, 2010; Weber *et al.*, 2014; Agez *et al.*, 2017; Cordshagen *et al.*, 2018), the dual (de)phosphorylation of threonine 906 and 1007 (Thr^906^/Thr^1007^) is a particularly potent regulator of KCC2 activity (Lee *et al.*, 2007, 2011; Rinehart *et al.*, 2009; Inoue *et al.*, 2012; Friedel *et al.*, 2015; Moore *et al.*, 2018). Thr^906^/Thr^1007^ becomes progressively dephosphorylated during neuronal development (Rinehart *et al.*, 2009; Friedel *et al.*, 2015). The phospho-mimetic threonine [T] to glutamate [E] (T906E/T1007E) mutations of KCC2 result in strong inhibition of the ion-transport activity in different cell lines and cultured neurons (Rinehart *et al.*, 2009; Inoue *et al.*, 2012; Friedel *et al.*, 2013). Unlike wild type KCC2 construct, who’s *in vitro* overexpression in immature neurons reduces [Cl^-^]_i_ and produces hyperpolarizing shift of GABA receptor type A (GABA_A_R) responses, the overexpression of T906E/T1007E mutant results in an increase in [Cl^-^]_i_, and a depolarizing shift of GABA_A_R responses (Inoue *et al.*, 2012; Friedel *et al.*, 2015).

The importance of precise post-translational KCC2 regulation has been corroborated by the identification of mutations that impair these processes in human patients with NDDs, including epilepsy (Kahle *et al.*, 2014; Merner *et al.*, 2015; Stödberg *et al.*, 2015). However, the pathophysiological consequences and mechanisms of impaired K-Cl co-transporters phosphorylation has only begun to be explored *in vivo* (Silayeva *et al.*, 2015, Kahle *et al.*, 2016*a*; Moore *et al.*, 2018).

Here, we have generated a KCC2 transgenic mouse carrying the phospho-mimetic threonine to glutamate [E] mutations at KCC2 Thr^906^ and Thr^1007^, which mimics constitutive phosphorylation and prevents normal developmental dephosphorylation at these sites (Rinehart *et al.*, 2009; Inoue *et al.*, 2012; Friedel *et al.*, 2015). *KCC2*^*E/E*^ homozygous mice die at birth, highlighting that precise phospho-regulation of these sites is essential for post-natal survival. In contrast, heterozygous (*KCC2*^*E/+*^) mice are viable and fertile. By studying these mice, we have found that heterozygous constitutive KCC2 Thr^906^/Thr^1007^ phosphorylation does not produce detectable changes in the overall abundance of KCC2, but significantly impairs the inhibitory strength of GABAergic neurotransmission and network properties during the first post-natal weeks of life. Moreover, *KCC2*^*E/+*^ mice exhibit increased seizure susceptibility and altered ultra-sonic vocalization at early ages (P2-P15) and social interaction deficits at adult ages (2 months). Postnatal treatment with the NKCC1 blocker bumetanide *in vivo* normalized the increased seizure susceptibility and aberrant network properties of *KCC2*^*E/+*^ mice at early ages but failed to restore social behavior at adult ages. Together, these results provide the first *in vivo* evidence of the importance of KCC2 post-translational modification for normal CNS development, and implicate dysregulated KCC2 Thr^906^/Thr^1007^ phosphorylation as a potential mechanism in the pathogenesis of NDDs.

## METERIALS AND METHODS

### Animals

The study was performed on *KCC2*^*E/+*^ and *KCC2*^*+/+*^ mixed background SV129/C57bl6-J mice. Animals were housed in a temperature-controlled environment with a 12-h light/dark cycle and free access to water and food. All procedures were in accordance with the European Communities Council Directive (86/609/EEC).

### Production of Kcc2 double point mutant targeted ES cell clones

Linearized targeting vector was transfected into 129Sv ES cells (genOway, Lyon, France) according to genOway’s electroporation procedures. PCR, Southern blot and sequence analysis of G-418 resistant ES clones revealed 2 clones as carrying the recombined locus. PCR over the 5’ end of the targeted locus was performed with a forward primer hybridizing upstream of the 5’ homology arm (5’- ATAGCGTTGGCTACCCGTGATATTGC-3’) and a reverse primer hybridizing within the Neomycin cassette (5’ AGGCTAGGCACAGGCTACATCCACAC-3’). Two Southern blot assays, assessing the correct recombination event at the 5’ end and at the 3’ end of the Kcc2 locus, were performed. These assays are based on the use of an internal and of an external probe, respectively. Finally, the integrity of the point mutations was confirmed by sequence analysis.

### Generation of chimeric mice and breeding scheme

Recombined ES cell clones were microinjected into C57BL/6 blastocysts, and gave rise to male chimeras with a significant ES cell contribution. These chimeras were bred with C57BL/6J mice expressing Cre-recombinase, to produce the Kcc2 double point mutant heterozygous line devoid of the Neomycin cassette. For each line, F1 genotyping was performed by PCR and Southern blot. PCR primers hybridizing upstream (5’- GTGGTTCGCCTATGGGATCTGCTACTC-3’)and downstream(5’-AGACAAGGGTTCATGTAACAGACTCGCC-3’) of the Neomycin cassette allowed identification of the Kcc2 endogenous, double point mutant allele harboring the Neomycin cassette, and double point mutant devoid of the Neomycin cassette (298-bp, 1946-bp and 387- bp, respectively). The Southern blot hybridized with an external probe allowed identification of the wild-type allele (14.1-kb) and the double point mutant allele (4.6-kb).

### Genotyping

DNA extraction was performed with KAPA Mouse Genotyping Kit. 88µl of water, 10µl of KAPA express extract buffer and 2µl of KAPA express extract enzyme was mixed and placed on the Eppendorf containing a small piece of biopsy. Lysis was performed on thermocycler during 10 min at 75°C for lysis and 5 min at 95°C for enzyme inactivation. The mix reaction for PCR was composed of 14.5µl H2O, 5µl 5X buffer promega, 1.5µl MgCl2 (25mM) promega, 1.25µl primers, 0.25µl dNTP (25mM), 0.25µl GoTaq polymerase promega pink for 1µl of DNA. The primers were following: 93598cof-KKA1; AGA CAA GGG TTC ATG TAA CAG ACT CGC C and 93599cof-KKA1; GTG GTT CGC CTA TGG GAT CTG CTA CTC.

### Hippocampal slice preparation and electrophysiological recordings

Brains were removed and immersed into ice-cold (2-4°C) artificial cerebrospinal fluid (ACSF) with the following composition (in mM): 126 NaCl, 3.5 KCl, 2 CaCl_2_, 1.3 MgCl_2_, 1.2 NaH_2_PO_4_, 25 NaHCO_3_ and 11 glucose, pH 7.4 equilibrated with 95% O_2_ and 5% CO_2_. Hippocampal slices (400 µm thick) were cut with a vibrating microtome (Leica VT 1000s, Germany) in ice cold oxygenated choline-replaced ACSF and were allowed to recover at least 90 min in ACSF at room (25°C) temperature. Slices were then transferred to a submerged recording chamber perfused with oxygenated (95% O_2_ and 5% CO_2_) ACSF (3 ml/min) at 34°C.

*Whole-cell patch clamp recordings* were performed from P15-P20 CA3 pyramidal neurons in voltage-clamp mode using an Axopatch 200B (Axon Instrument, USA). To record the spontaneous synaptic activity, the glass recording electrodes (4-7 MΩ) were filled with a solution containing (in mM): 100 KGluconate, 13 KCl, 10 HEPES, 1.1 EGTA, 0.1 CaCl_2_, 4 MgATP and 0.3 NaGTP. The pH of the intracellular solution was adjusted to 7.2 and the osmolality to 280 mOsmol l^-1^. The access resistance ranged between 15 to 30 MΩ. With this solution, the GABA_A_ receptor-mediated postsynaptic current (GABAA-PSCs) reversed at - 70mV. GABA-PSCs and glutamate mediated synaptic current (Glut-PSCs) were recorded at a holding potential of −45mV. At this potential GABA-PSC are outwards and Glut-PSCs are inwards. All recordings were performed using Axoscope software version 8.1 (Axon Instruments) and analyzed offline with Mini Analysis Program version 6.0 (Synaptosoft). For the acute bumetanide treatment, before recording, slices were incubated during 3h in ACSF containing 10µM of bumetanide.

*Single GABA*_*A*_ *and NMDA channel recordings* were performed from P7 to P30 visually identified hippocampal CA3 pyramidal cells in cell-attached configuration using Axopatch-200A amplifier and pCLAMP acquisition software (Axon Instruments, Union City, CA). Data were low-pass filtered at 2 kHz and acquired at 10 kHz. The glass recording electrodes (4-7 MΩ) were filled with a solution containing (in mM): (1) for recordings of single GABA_A_ channels: GABA 0.01, NaCl 120, KCl 5, TEA-Cl 20, 4-aminopyridine 5, CaCl_2_ 0.1, MgCl_2_10, glucose 10, Hepes-NaOH 10 (Tyzio *et al.*, 2003, 2014); (2) for recordings of single NMDA channels: nominally Mg^2+^ free ACSF with NMDA (10 µM) and glycine (1 µM) (Tyzio *et al.*, 2003). The pH of pipette solutions was adjusted to 7.2 and the osmolality to 280 mOsmol l^-1^. Both single GABA_A_ and single NMDA channel currents were recorded in voltage-clump mode at different membrane potentials (from −80mV to 80mV for GABA_A_ and from −120 to 40 mV for NMDA) in order to visualize outwardly and inwardly directed single channel currents. Experiments with recorded only outward or only inward currents were excluded from analysis. Analysis of currents trough single channels and I-V curves were performed using Clampfit 9.2 (Axon Instruments) as described by (Tyzio *et al.*, 2003).

*Extracellular recording of the local field potentials (LFPs)* were performed from P5-P20 CA3 region of hippocampus. Extracellular 50µm tungsten electrodes (California Fine Wire) were placed in the pyramidal cell layer to record the Multi Unit Activity (MUA). The signals were amplified using a DAM80i amplifier, digitized with an Axon Digidata 1550B, recorded with Axoscope software version 8.1 (Axon instruments) and analyzed offline with Mini Analysis Program version 6.0 (Synaptosoft). The frequency MUA was analyzed before, during and after 2 min of application of isoguvacine (10µM). The effect of drug application was determined by the percentage of frequency changes between the frequency obtained during the application of the drug compared to the frequency obtained before the application. The changes >20% were considered as excitation. Conversely, the changes <20% were considered as inhibition. Slices were classified as no responses when the changes of frequency were lower than 20%.

### Antibodies for Western blot

The following antibodies were raised in sheep and affinity-purified on the appropriate antigen by the Division of Signal Transduction Therapy Unit at the University of Dundee: KCC2A phospho-Thr 906 (SAYTYER(T)LMMEQRSRR [residues 975 - 989 of human KCC3A] corresponding to SAYTYEK(T)LVMEQRSQI [residues 899 - 915 of human KCC2A]) (Catalog number: S959C); KCC2A phospho-Thr 1007 (CYQEKVHM(T)WTKDKYM [residues 1032 - 1046 of human KCC3A] corresponding to TDPEKVHL(T)WTKDKSVA [residues 998 - 1014 of human KCC2A]) (Catalog number: S961C); KCC2 phospho-Ser940 (Catalog number: NBP2-29513). Pan KCC2 total antibody (residues 932-1043 of human KCC2) was purchased from NeuroMab (Catalog number: 73-013). Anti-β-Tubulin III (neuronal) antibody was purchased from Sigma-Aldrich (Catalog number: T8578). Secondary antibodies coupled to horseradish peroxidase used for immunoblotting were obtained from Pierce. IgG used in control immunoprecipitation experiments was affinity-purified from pre-immune serum using Protein G-Sepharose.

### Buffers for Western Blots

Buffer A contained 50 mM Tris/HCl, pH7.5 and 0.1mM EGTA. Lysis buffer was 50 mM Tris/HCl, pH 7.5, 1 mM EGTA, 1 mM EDTA, 50 mM sodium fluoride, 5 mM sodium pyrophosphate, 1 mM sodium orthovanadate, 1% (w/v) Triton-100, 0.27 M sucrose, 0.1% (v/v) 2-mercaptoethanol, and protease inhibitors (complete protease inhibitor cocktail tablets, Roche, 1 tablet per 50 mL). TBS-Tween buffer (TTBS) was Tris/HCl, pH 7.5, 0.15 M NaCl and 0.2% (v/v) Tween-20. SDS sample buffer was 1X NuPAGE LDS sample buffer (Invitrogen), containing 1% (v/v) 2-mercaptoethanol. Protein concentrations were determined following centrifugation of the lysate at 16,000 × g at 4°C for 20 minutes using the Bradford method with bovine serum albumin as the standard.

### Immunoprecipitation with phosphorylation site-specific antibodies

KCCs phosphorylated at the KCC2 Thr^906^ and Thr^1007^ equivalent residue were immunoprecipitated from clarified hippocampal and cortical culture lysates (centrifuged at 16,000 x g at 4°C for 20 minutes) using phosphorylation site-specific antibody coupled to protein G–Sepharose as described (Los Heros *et al.*, 2014). The phosphorylation site-specific antibody was coupled with protein-G–Sepharose at a ratio of 1 mg of antibody per 1 mL of beads in the presence of 20 µg/mL of lysate to which the corresponding non-phosphorylated peptide had been added. 2 mg of clarified cell lysate were incubated with 15 µg of antibody conjugated to 15 µL of protein-G–Sepharosefor 2 hours at 4°C with gentle agitation. Beads were washed three times with 1 mL of lysis buffer containing 0.15 M NaCl and twice with 1 mL of buffer A. Bound proteins were eluted with 1X LDS sample buffer.

### Immunoblotting

Hippocampi tissue lysates (15 µg) in SDS sample buffer were subjected to electrophoresis on polyacrylamide gels and transferred to nitrocellulose membranes. The membranes were incubated for 30 min with TTBS containing 5% (w/v) skim milk. The membranes were then immunoblotted in 5% (w/v) skim milk in TTBS with the indicated primary antibodies overnight at 4°C. Antibodies prepared in sheep were used at a concentration of 1-2 µg/ml. The incubation with phosphorylation site-specific sheep antibodies was performed with the addition of 10 µg/mL of the nonphosphorylated peptide antigen used to raise the antibody. The blots were then washed six times with TTBS and incubated for 1 hour at room temperature with secondary HRP-conjugated antibodies diluted 5000-fold in 5% (w/v) skim milk in TTBS. After repeating the washing steps, the signal was detected with the enhanced chemiluminescence reagent. Immunoblots were developed using a film automatic processor (SRX-101; Konica Minolta Medical) and films were scanned with a 600-dpi resolution on a scanner (PowerLook 1000; UMAX). Figures were generated using Photoshop and Illustrator (Adobe). The relative intensities of immunoblot bands were determined by densitometry with ImageJ software.

### Behavior

All behavioral procedures were carried between 8:00 and 17:00 h under dim light conditions. Male animals were moved to the testing room in their home cages 30 min prior to test beginning. The same animals have been used for all adulthood behavioral tests, starting at 6- week old. All experiments and analysis were done in blind genotyping by only one experimenter.

#### Vocalization

To induce vocalization, pups were isolated individually from their mother at P2-4-8-10-12. They were placed into an isolation box (23×28×18 cm) located inside a sound attenuating isolation cubicle (54×57×41 cm; Coulbourn Instruments, Allentown, PA, USA).

An ultrasound microphone sensitive to frequencies of 10 to 250 kHz (Avisoft UltraSoundGate Condenser microphone capsule CM16/CMPA, Avisoft Bioacoustics, Berlin, Germany) was placed in the roof of the box. Vocalizations were recorded for 3 minutes using the Avisoft Recorder software (version 4.2) with a sampling rate of 250 kHz in 16 bit format. Recordings were transferred to SASLab Pro (version 5.2; AvisoftBioacoustics) and a fast Fourier transformation was conducted (512 FFT-length, 100% frame, Hamming window and 75%time window overlap, cut off frequencies high pass 20Khz) before analyzing the number of calls emitted by mice.

#### Open field

The mice were individually placed in the square field (38.5 x 38.5cm; Noldus, Netherlands). Recording began 10 seconds after the mouse was placed inside the apparatus for a 10 minutes trial. Behaviors were recorded by a video camera fixed above the apparatus and analyzed using the Ethovision 11.5 software (Noldus, Netherlands). After each trial, the open field was cleaned with a solution containing 70% of ethanol. Anxiety and locomotor-like behaviors were analyzed using three parameters: the time spent in the center (12.8cm X 12.8cm), the number of entries in the center and the distance travelled.

#### Social interaction

The 3-chamber test was performed as previously described (Peça *et al.*, 2011). The test was conducted in an apparatus (59 x 39.5cm; Noldus, Netherlands) divided in a central empty compartment (19.5 x 39.5 cm) and two side-compartments (19.5 x 39.5 cm) containing a plastic cup-like cage for the strangers. The strangers were of the same sex and age as the tested mice and were habituated during 15min (one habituation per day) to the plastic cup-like cage 4 days prior to the beginning of the test. The mouse was recorded during 4 consecutive trials of 5 min. Trial 1, habituation: the tested mouse was placed in the empty compartment with the access to the other compartments closed. Trial 2, sociability testing: a first stranger was placed in a plastic cup-like cage in one of the two-side compartments and the tested mouse could explore freely all the compartments. Trial 3, post-test: the tested mouse could explore freely its environment and be habituated for a longer time to the stranger 1 (familiar stranger). Trial 4, social novelty testing: the second stranger (novel stranger) was placed in the second plastic cup-like cage in the opposite side-compartment. For each tested mouse, the side-compartments where the strangers were placed were alternated to avoid any side preferences. All trials were recorded by a video camera placed above the apparatus by using the Ethovision 11.5 software (Noldus, Netherlands) and the time spent in each chamber was manually analyzed. Trials where the mice have returned less than once time to the chamber are removed.

#### Splash

The splash test was performed as previously described (Moretti et al; 2015). Mice were sprayed in the dorsal coat with a 10% sucrose solution. The viscosity of this solution dirties the mice and initiates the grooming behavior. After being sprayed, mice were individually placed in a plexiglass cylinder (15 × 45cm; Form X.L., France) and their behaviors were recorded for 5 minutes. Self-care and motivational behaviors were manually analyzed using three parameters: the duration of grooming, the latency to start the first grooming, and the frequency of grooming events. A grooming event was defined as at least one episode of any category of grooming (paw licking, head wash, body groom, leg licking, tail/genital licking).

#### Seizure testing

For testing of seizure susceptibility, we used convulsant agent flurothyl (2,2,2-trifluoroethyl ether, Sigma) that is widely used to study the epilepsy in different animal models (Villeneuve *et al.*, 2000; Velíšek *et al.*, 2011). The advantage of the flurothyl, as compared to other pro-convulsive agents is that it could be used to induce epilepsy-like activity in juvenile P0-P30 rats and mice (Villeneuve *et al.*, 2000; Velíšková *et al.*, 2017). Because the latencies of the effects of flurothyl depend on large number of parameters including atmospheric pressure, humidity, temperature, movements of the air in experimental chamber, animal weight and age (Velíšková *et al.*, 2017) all experiments were performed using pairs of littermate males (P15 and P30) that were placed in transparent hermetic two-compartment cage inhalated with flurothyl. The progressive injection of the flurothyl into the cage produced a stereotypical behavioral manifestation of limbic seizure episodes that varied on dependence of the animal age and genotype. At P15 these episodes commenced in all studied animals with forelimb and/or tail extension, rigid posture (stage 1) followed by subconvulsions (stage 2), one to three brief (1-2s) myoclonic jerks (stage 3), severe tonic-clonic seizures (stage 4) lasting 5-20 s that ended with falling and immobility of the animal (stage 5) (**Fig. 4A**). At P30 seizure episodes started from animal immobility and ended by severe tonic-clonic seizures (stage 4). The intermediate stages (rigid posture, subconvulsions) were absent or not clearly detectable at this age. 10 s after beginning of tonic-clonic seizures in more resistant animal from each pair, the injection of flurothyl was discontinued and cage was inhalated with fresh air. 2-5 min after stopping of the exposure to flurothyl most of mice returned to their 4-limb horizontal position, but remained immobile during 5-10 min. Thereafter all mice started moving and exploring the cage, although their moving activity was not scored.

**Figure 1.**
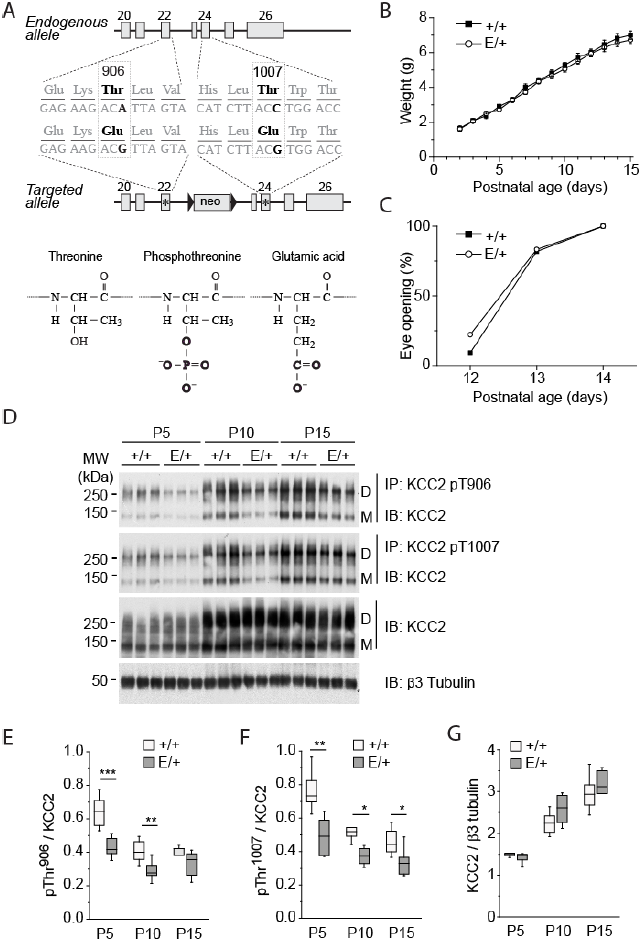
Characterization of *KCC2*^*E/+*^ phospho-mimetic mice. **A)** Mutagenesis scheme. Thr^906^ and Thr^1007^ are located in exons 22 and 24, respectively, and are preserved in endogenous allele. T906E and T1007E mutations are located in the second targeted allele that contains a Neomycin selection cassette excised *in vivo* by Cre recombinase. **B)** Weight gain by *KCC2*^*E/+*^ and *KCC2*^*+/+*^pups. The plot represents mean±SEM of values obtained from 11 *KCC2*^*+/+*^ pups (4 litters) and 18 *KCC2*^*E/+*^ pups (4 litters). Two-way Anova analysis revealed no statistically significant difference between *KCC2*^*E/+*^ and *KCC2*^*+/+*^pups (*P*=0.18, F=1.84) and strong difference of weight during development (*P*=1.1E-8, F=205.39). **C)** Age dependence of eye opening, an external sign of normal CNS maturation in vertebrates (Stromland and Pinazo-Durán, 2002), in *KCC2*^*E/+*^ and *KCC2*^*+/+*^pups. The plot represents percentage of animals with open eyes (n=11 *KCC2*^*+/+*^ pups [4 litters] and 18 *KCC2*^*E/+*^ pups [4 litters]). The chi-square analysis revealed no statistically significant difference between *KCC2*^*E/+*^ and *KCC2*^*+/+*^pups (*P*=0.83472). **D)** Abundance of KCC2 in *KCC2*^*E/+*^ and *KCC2*^*+/+*^pups. Hippocampal lysates from *KCC2*^*E/+*^ and *KCC2*^*+/+*^ at the indicated time pointspoints were subjected to immuno-precipitation (IP) with the indicated phosphorylation site-specific antibodies recognizing pThr906- or pThr1007-KCC2, and the immuno-precipitated products were detected with the pan-KCC2 antibody (IB). The same antibody was used for detection of total KCC2 protein abundance in the same input material. An antibody recognizing neuron specific ²3 tubulin was employed to normalize total protein amounts for sample loading. **E,F,G)** Developmental abundance of immunoprecipitated phosphorylated forms of Thr^906^ (E), Thr^1007^ (E) and total KCC2 (G). Data are from n=6 per condition from 4 different litters. * *P* <0.05; ** *P* <0.01. For more details on statistical tests see **Table S1.**

**Figure 2.**
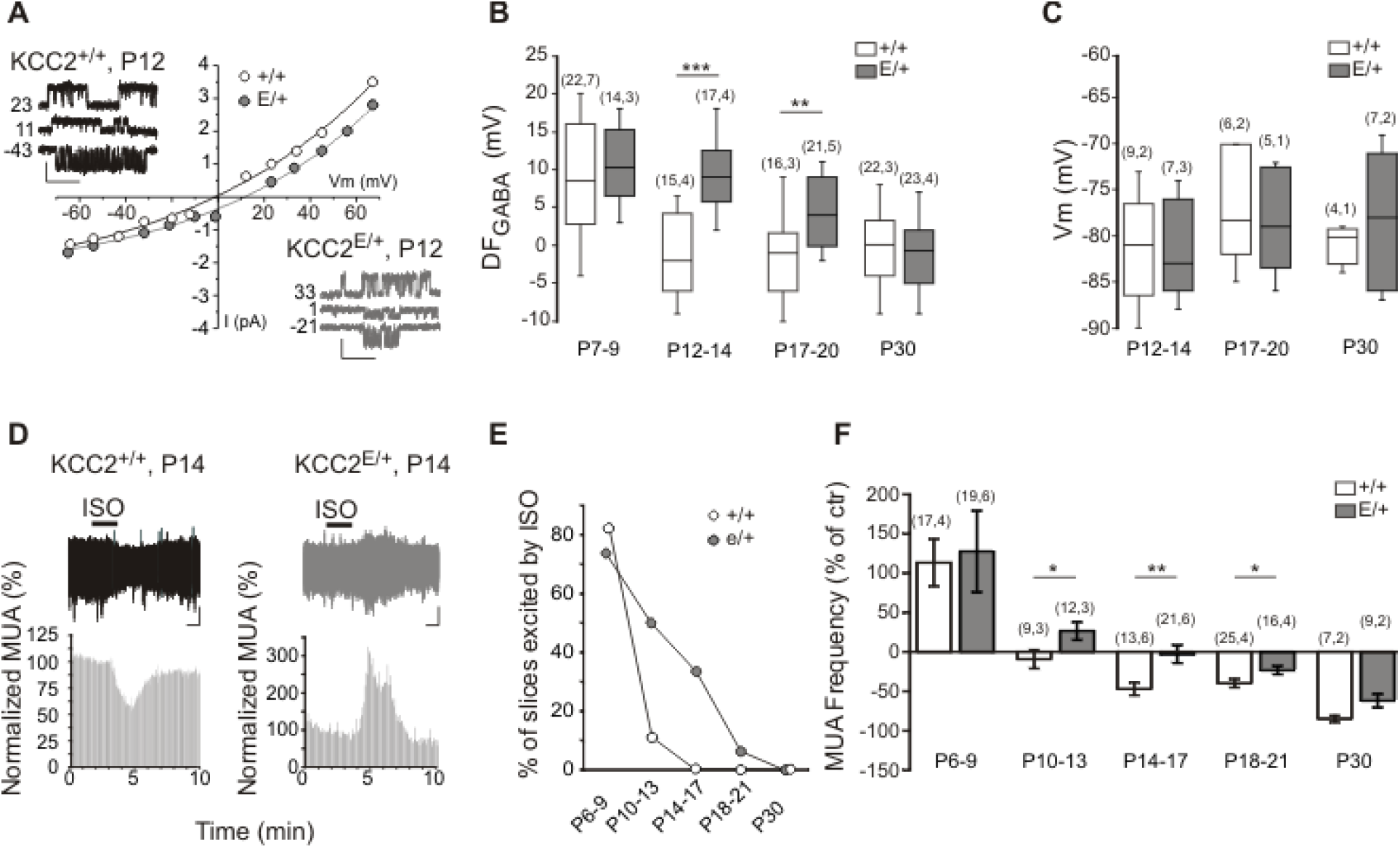
Delayed depolarizing shift of GABA transmission in *KCC2*^*E/+*^ CA3 neurons. A) Representative example of I-V curves and traces at different membrane potentials of single GABA_A_ channel currents recorded from CA3 neurons at P12. DF_GABA_ was determined as the intercept of the I-V curve with the X-axis. Scale values: 1 s, 1 pA. **B)** Boxplots of DF_GABA_ at different age frames. Numbers in parenthesis indicate the number of recorded neurons (first value) and number of animals (value after coma). **p<0.01; ***p<0.001. **C)** Boxplots of resting membrane potential (Vm) in CA3 neurons of indicated age frames analyzed using single NMDA-channel recording as shown in **Fig. S2D)** Traces illustrate the representative field potential recordings from CA3 neurons of inhibitory (left plot) and excitatory (right plot) actions of isoguvacine applied as indicated with horizontal bar. Scale values: 1 min, 20 µV. The histograms show the quantification of spike frequencies in illustrated traces. **E)** Percentage of slices showing an increase of MUA in response to isoguvacine. **F)** Mean ± SEM of the relative change of isoguvacine-dependent MUA frequency at different age frames. Numbers in parenthesis indicate the number of recorded slices (first value) and number of animals (value after coma). *p<0.05; **p<0.01. For more details on statistical tests see **Table S2.**

**Figure 3.**
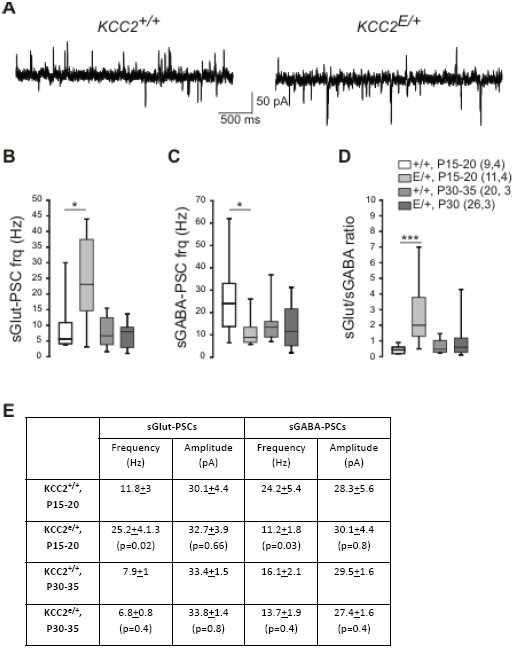
The *KCC2*^*E/+*^ P15 but not P30 mice show an altered glutamate/GABA balance. A) Representative whole cell recordings of spontaneous post-synaptic currents (holding potential =-45mV) in 2 week-old *KCC2*^*+/+*^ and *KCC2*^*E/+*^ CA3 pyramidal neurons. Scale values: 500 ms, 50 pA. (B-C) Boxplot of the frequency of the spontaneous glutamatergic (B) and GABAergic (C) post-synaptic currents recorded from 2 and 4 week-old *KCC2*^*+/+*^ and *KCC2*^*E/+*^ CA3 pyramidal neurons. (D) Boxplots of the frequency ratio of spontaneous glutamatergic to GABAergic postsynaptic currents. Numbers in parenthesis indicate the number of cells recorded and mice used. The mean±SEM age of recorded neurons was identical for 2 week-old *KCC2*^*E/+*^ and *KCC2*^*+/+*^ mice (17±0.8 and 17±0.7, respectively). *P<0.05, ***P<0.01, one-way Anova test.

**Figure 4.**
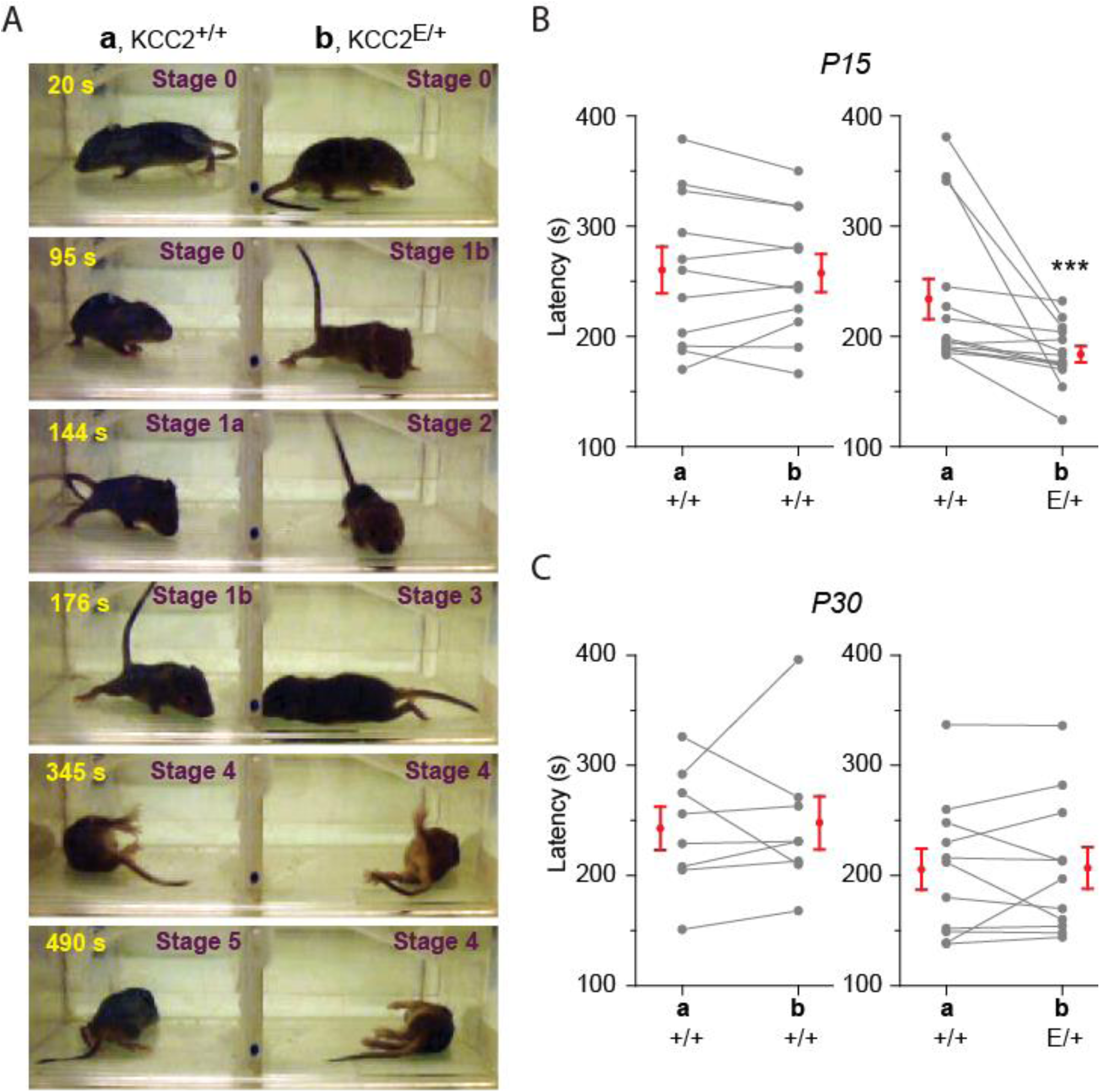
*KCC2*^*E/+*^ P15 but not P30 mice show an increased onset of flurothyl-induced seizures. **A)** Photos illustrating different stages of P15 mice responses to flurothyl. The first behavioral response is animal immobility (Stage 0) followed by rigid posture (Stage 1a) and tail extension (Stage 1b). The stages 0 and 1 were observed in 100% of P15 mice. After Stage 1a, ∼50% of animals exhibited brief (1-2s) subconvulsive events (Stage 2) and/or 1^st^ short seizure (1-2s) episodes (Stage 3) followed by severe long-lasting (10-30 s) generalized (tonic-clonic) seizures (Stages 4). After tonic-clonic seziures, mice returned to four-limb posture and remain immobile for 10-30 min (Stage 5). **B, C)** Latency of onset of tonic-clonic seizures i.e. latency to stage 4 of P15 (B) and P30 (C) *KCC2*^*+/+*^ and *KCC2*^*E/+*^ mice placed in compartments **a** and **b** as indicated. Connected points indicate pairs of animals. The points with error bars indicate mean ± SEM values.

Flurothyl was progressively injected into the cage using nano-pump (Harvard apparatus) and homogeneously distributed using mini-ventilator incorporated into the chamber. Behavioral responses were recorded using a video camera. The latency of tonic-clonic seizures was determined post-hoc as time of the first body convulsion in continuous series of tonic-clonic seizures.

#### Statistical analysis

Statistical analyses were conducted with OriginPro 9.0.0 which also indicated that assumptions of normality (Shapiro-Wilk test) and equal variance (Brown-Forsyth test) were met. A *P*<0.05 was considered significant for these and all subsequent tests. For data displaying normal distribution and equal variance one-way or two-way (as indicated) ANOVA and the post hoc Tukey test were used for multiple comparisons between groups. For data displaying non-normal distribution or unequal variance, Mann–Whitney U-test was used for comparison between 2 independent groups and Wilcoxon matched pairs test was employed to compare paired data. The Chi-square test was used to determine the significance of the difference between the frequencies of event occurrence.. Sample size for each group reported in electrophysiological experiments is at least 5 neurons from at least 3 independent experiments. For the boxplots, the box extends from the first (Q1) to third (Q3) quartiles. The line inside the box represents the median. The whiskers define the outermost data point that falls within upper inner and lower inner fence (Q1-1.5(IQR)) and (Q3-1.5(IQR)), respectively.

## RESULTS

### Characterization of *KCC2*^*E/+*^ mice

To determine the *in vivo* significance of phosphorylation at the WNK/SPAK-kinase dependent KCC2 Thr^906^/Thr^1007^ phosphorylation motif (Rinehart *et al.*, 2009; Inoue *et al.*, 2012; Friedel *et al.*, 2015), we generated mice harboring phospho-mimetic glutamic acid substitutions at these sites (T906E/T1007E) via homologous recombination (**Fig. 1A**). Homozygous *KCC2*^*E/E*^ mice died within the first 4-12 hours after birth from apparent respiratory distress, but heterozygous *KCC2*^*E/+*^ mice were viable, fertile and survived through adulthood. Compared to *KCC2*^*+/+*^ littermates, the *KCC2*^*E/+*^ mice showed no difference in the time of eye opening or in weight gain from postnatal days (P) 0 to P15 (**Fig. 1B and C**). Immunoprecipitation of KCC2 from hippocampal lysates using antibodies that recognize the native phosphorylated, but not mutated (T906E and T1007E) forms of KCC2 (Los Heros *et al.*, 2014), showed a significant decrease in *KCC2*^*E/+*^ mice as compared to *KCC2*^*+/+*^ littermates (**Fig. 1D,E and F**, two way-Anova test, see **Table S1** for details on statistic). The abundance of total KCC2 in hippocampal lysates from *KCC2*^*+/+*^ mice increased progressively from P5 to P15 (**Fig. 1D and G**, *P*=3.7E-14, two-way Anova, n=6 per condition from 4 different litters [n=6, 4]), in agreement with previous reports (Lu *et al.*, 1999; Stein *et al.*, 2004). A similar developmental up-regulation of KCC2 abundance was observed in hippocampi from *KCC2*^*E/+*^ mice (**Fig. 1G**). No difference in total KCC2 abundance was seen between *KCC2*^*+/+*^ and *KCC2*^*E/+*^ mice at all studied time points (**Fig. 1D and G**, *P*=0.25, two-way Anova, [n=6, 4]), or KCC2 Ser^940^ phosphorylation (**Fig. S1A and B**, *P*=0.69, two-way Anova, [n=6, 4]). In addition, no difference in the total abundance of NKCC1 or the NKCC1/KCC2 regulatory kinases WNK1, WNK3, SPAK, or OSR1 were noted in *KCC2*^*+/+*^ and *KCC2*^*E/+*^ mice (**Fig. S1C and D, Table S1**).

### *KCC2*^*E/+*^ CA3 pyramidal neurons exhibit a delay in the GABAergic developmental sequence

Phospho-mimetic KCC2 T906E/T1007E mutation impairs Cl^-^ extrusion capacity *in vitro* (Inoue *et al.*, 2012; Friedel *et al.*, 2015). To test whether *in vivo* T906E/T1007E mutations alter neuronal Cl^-^ homeostasis, we measured the driving force of GABA_A_ channels (DF_GABA_) in CA3 pyramidal neurons of acute hippocampal slices from *KCC2*^*+/+*^ and *KCC2*^*E/+*^ mice using non-invasive cell-attached single-GABA_A_ channel recordings (Tyzio *et al.*, 2003, 2014). In agreement with previous reports (Tyzio *et al.*, 2006, 2014), in slices prepared from P7-P9 *KCC2*^*+/+*^ mice, the DF_GABA_ was positive in most recorded neurons (**Fig. 2B**, DF_GABA_ =8.5±1.5 mV, (22 neurons, 7 mice [n=22,7])). Onwards from P12-14, DF_GABA_ shifted towards less depolarizing and even hyperpolarizing values (**Fig. 2A and B**, DF_GABA_=-1.4±1.3 mV, [n=15, 4], *P*=1.26E-5, post-hoc Tukey test). In contrast, in CA3 pyramidal neurons from *KCC2*^*E/+*^ mice, DF_GABA_ remained strongly depolarized until P17-20 (**Fig. 2A and B**, DF_GABA_=9.4±1.1 mV [n=17, 4]), and was significantly different compared to age matched neurons from *KCC2*^*+/+*^ mice (*P*=5.7E-7 and *P*=0.002 for P12-14 and P17-20 age frames, respectively, two-way Anova and post-hoc Tukey test, **Table S2**). Interestingly, the hyperpolarizing shift in *KCC2*^*E/+*^ mice occurred at P30 (**Fig. 2B**, DF_GABA_=-1.2±1.0 mV [n=23,4], *P*=0.03 compared to *KCC2*^*E/+*^ P17-20 values, *P*=0.5 compared to *KCC2*^*+/+*^ age matched, post-hoc Tukey tests). There was, however, no significant difference in the resting membrane potential of *KCC2*^*E/+*^ and *KCC2*^*+/+*^ CA3 pyramidal neurons at P12-14 and P17-20 (**Fig. 2C, Fig. S2 and Table S2**). These data show that the developmental hyperpolarizing GABA shift is delayed in the CA3 pyramidal neurons of *KCC2*^*E/+*^ mice.

We next examined the emergence of functional GABAergic inhibition in the developing *KCC2*^*+/+*^ and *KCC2*^*E/+*^ hippocampus. We performed non-invasive extracellular recordings of MUA (Khazipov *et al.*, 2004; Ben-Ari *et al.*, 2007) in acute hippocampal slices from *KCC*^*+/+*^ and *KCC2*^*E/+*^ littermates, and investigated the effect of bath application of the GABA_A_R agonist isoguvacine (10µM) on the firing of the CA3 pyramidal neurons from P6 to P30. Consistent with previous results (Ben-Ari *et al.*, 2007), in wild-type *KCC2*^*+/+*^ mice, isoguvacine induced an increase of firing of CA3 pyramidal neurons in 83% of P6-9 slices (**Fig. 2E**, 17 slices, 4 mice [n=17, 4]). Starting from P14-17 no firing increase was observed in response to isoguvacine (**Fig. 2D and E**). Remarkably, in *KCC2*^*E/+*^ mice, the isoguvacine-induced increase of firing persisted until P18 (**Fig. 2D and E**, *P*=1E-5, Chi-square test). Averaging the overall effect of isoguvacine on the firing of CA3 pyramidal neurons confirmed the delayed hyperpolarizing shift of GABA action in *KCC2*^*E/+*^ mice (**Fig. 2F, Table S2**).

### Altered glutamate/GABA balance in juvenile *KCC2*^*E/+*^ CA3 pyramidal neurons

A loss of equilibrium in glutamatergic and GABAergic synaptic drive has been observed in several NDDs associated with a delayed developmental GABA shift (Tyzio *et al.*, 2014; Deidda *et al.*, 2015; Banerjee *et al.*, 2016). We therefore performed whole cell recordings of CA3 pyramidal neurons in acute hippocampal slices at P15-20 and P30 to measure spontaneous GABA_A_R-mediated and glutamate mediated postsynaptic currents (sGABA-PSCs and sGlut-PSCs) with a low Cl^-^ pipette solution (**Fig. 3A-D)**. At P15-20, we found that the frequency of sGlut-PSCs, but not their amplitude, was increased in *KCC2*^*E/+*^ CA3 pyramidal neurons compared to *KCC2*^*+/+*^ littermates (**Fig. 3A-E**). In contrast, the frequencies of sGABA-PSCs were decreased in *KCC2*^*E/+*^ CA3 pyramidal neurons, with no change in amplitude (**Fig. 3A-E**). At P30, whole cell recordings revealed no significant difference in the frequency or amplitude of sGlut-PSCs and sGABA-PSCs (**Fig. 3B-E**). Calculation of the ratio of sGlut-PSCs and sGABA-PSCs frequencies (hereafter referred to as excitatory-inhibitory (E/I) balance) recorded from the same CA3 pyramidal neurons showed a significant increase in *KCC2*^*E/+*^ neurons compared to *KCC2*^*+/+*^ neurons (2.71±0.6 in *KCC2*^*E/+*^ vs 0.44+0.09 in *KCC2*^*+/+*^, *P*=0.005, one-way Anova test, **Fig. 3D, E**). There was, however, no difference in the E/I balance at P30-35 (0.61±0.09 in *KCC2*^*E/+*^ vs 0.97±0.2 in *KCC2*^*+/+*^, *P*=0.4, one-way Anova test, **Fig. 3D, E**). These data show that the E/I balance is impaired in CA3 neurons of P15-P20 *KCC2*^*E/+*^ mice, coinciding with the depolarizing DF_GABA_.

### Increased seizure susceptibility in juvenile *KCC2*^*E/+*^ mice

Genetic studies have shown an association between KCC2 dysfunction and different types of human epilepsy syndromes (Kahle *et al.*, 2016*b*; Moore *et al.*, 2017). To determine whether impaired KCC2 Thr^906^/Thr^1007^ phosphorylation impacts seizure susceptibility, pairs of immature and juvenile (P15 and P30) *KCC2*^*+/+*^ or *KCC2*^*E/+*^ mice were placed in a two-compartment hermetic chamber (thereafter termed compartments **a** and **b**), thereby ensuring simultaneous exposure of both animal groups to the convulsant agent flurothyl (2,2,2- trifluoroethyl ether) (see Methods). Flurothyl induced severe tonic-clonic seizures with a loss of posture and jumping in both *KCC2*^*+/+*^ and *KCC2*^*E/+*^ mice (stage 4 in **Fig. 4A**). As a control, when both compartments included P15 wild-type *KCC2*^*+/+*^ animals, the latencies of seizure appearance in both animals were similar (**Fig. 4B**, *P*=0.37, Wilcoxon Paired test, n=11 pairs). When compartment **a** included P15 wild-type *KCC2*^*+/+*^ mice and compartment **b** contained their heterozygous *KCC2*^*E/+*^ littermates, the seizure latency of heterozygous *KCC2*^*E/+*^ mice was ∼20 % shorter as compared to their wild-type *KCC2*^*+/+*^ littermates (**Fig. 4B**, 234±18s vs 184±7s, *P*=2.4E-4, Wilcoxon Paired test, n=14 pairs). In contrast, 30 day-old *KCC2*^*+/+*^ and *KCC2*^*E/+*^ mice had no difference in seizure latency (**Fig. 4C**, *P*=0.92, Wilcoxon Paired test, n=11 pairs). These results show *KCC2*^*E/+*^ mice have increased seizure susceptibility at P15, coinciding with their depolarizing DF_GABA_ and increased neuronal network activity.

### Altered vocalization and social interactions in *KCC2*^*E/+*^ mice

An increase in seizure susceptibility is a co-morbidity frequently associated with NDDs (Agrawal and Govender, 2011). In addition, similar to described above changes in DF_GABA_ and alterations in synaptic activity were reported in different animal models of NDD (Tyzio *et al.*, 2014; Deidda *et al.*, 2015; Banerjee *et al.*, 2016). We therefore assessed several neuro-behavioral tests in *KCC2*^*+/+*^ and *KCC2*^*E/+*^ mice, including ultrasonic vocalization (USVs) (alteration of communication) (Scattoni *et al.*, 2009), open field (anxiety and hyperactivity) (Belzung and Griebel, 2001), three-chamber test (sociability) (Silverman *et al.*, 2010), and grooming splash tests (depression) (Wang *et al.*, 2018).

USVs test were assessed in P2 to P12 mice. In *KCC2*^*+/+*^ mice, the number of calls varied at different postnatal days and showed a typical ontogenetic profile (Wiaderkiewicz *et al.*, 2013) (**Fig. 5A**). *KCC2*^*E/+*^pups compared to *KCC2*^*+/+*^ animals showed a statistically significant different ontogenetic profile (**Fig. 5A**; *P*=0.003, two-way Anova test). In P2 to P8 *KCC2*^*E/+*^ mice, the number of ultrasonic calls was similar to *KCC2*^*+/+*^ animals but at P10 and P12 the number of calls was higher (**Fig. 5A and B**; *P*=0.005 and *P*=0.003 for P10 and P12, respectively, post-hoc Tukey test).

**Figure 5.**
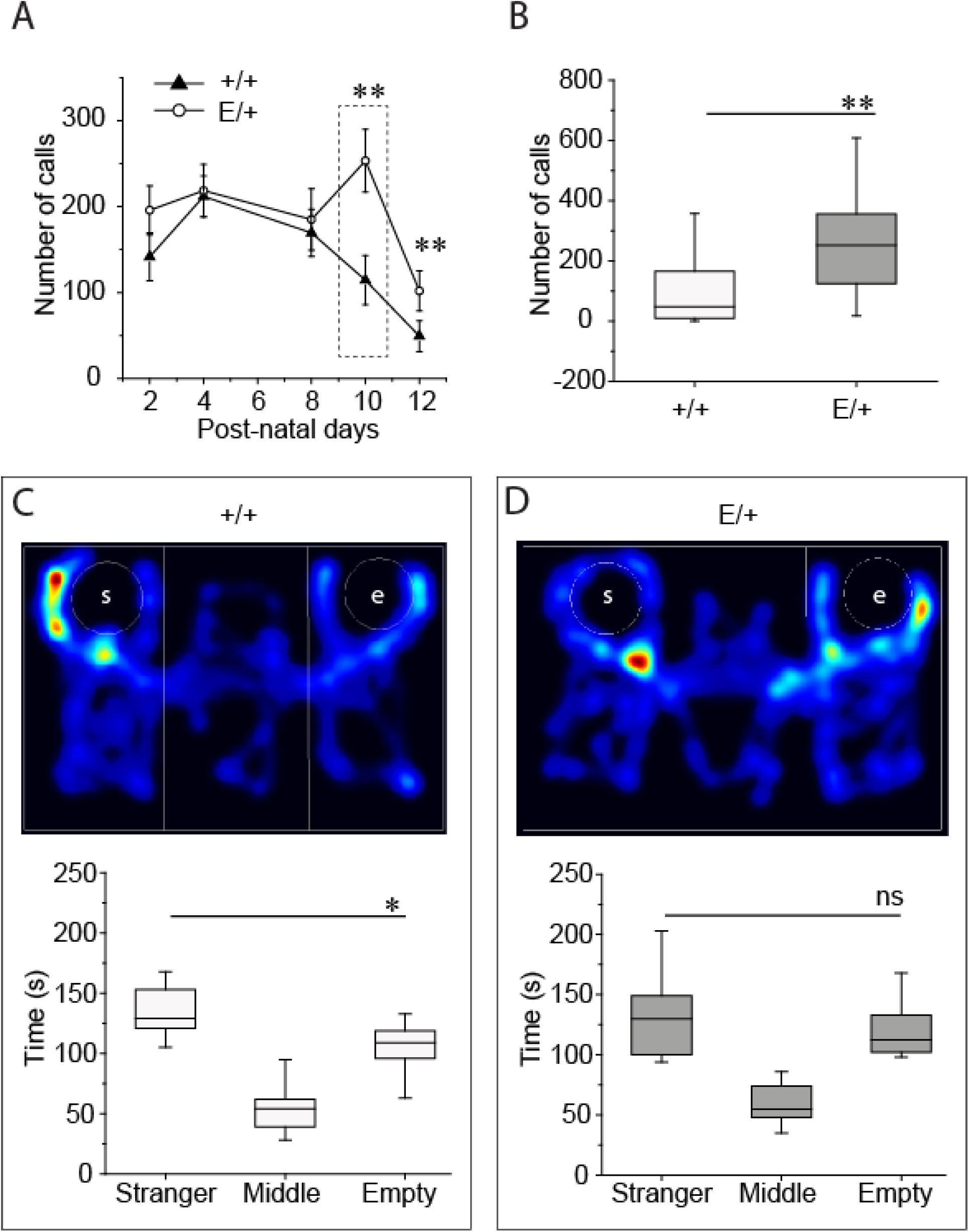
KCC2 T906E/T1007E impairs newborn communication and adult social-behavior. **A)** Number of calls in responses to separation from mother during 3 min sessions at P2-P12. Data are expressed as mean ± SEM of calls. **B)** Boxplot of number of calls emitted by WT and E/+ mice at P10. Triangles (*KCC2*^*+/+*^) and circles (*KCC2*^*E/+*^) show the distribution of measurements. **C-D)** Time spent by *KCC2*^*+/+*^ (C) and *KCC2*^*E/+*^ (D) mice in stranger chamber and empty chamber. Upper panels illustrate heat maps of mice movements in the sociability tests. White circles show location of plastic cup-like cage containing stranger (s) or empty (e). Boxplots illustrate the quantification of the time spent by tested mice in indicated compartments. ns= non-significant, \**P*<0.05.

The three-chamber test that was performed on P60 mice using widely explored paradigm (Peça *et al.*, 2011) revealed difference in social behavior of *KCC2*^*+/+*^ and *KCC2*^*E/+*^ mice; while the wild type *KCC2*^*+/+*^ mice spent ∼35% more time to explore the compartment with stranger than the empty compartment (**Fig. 5C**, *P*=3.7E-4, one-way Anova test), the *KCC2*^*E/+*^ mice showed no interest to stranger and spent similar times in empty and stranger-containing compartments (**Fig. 5D**, *P*=0.1, one-way Anova test). The three-chamber test allows determination of social novelty and exploration behavior. Both *KCC2*^*+/+*^ and *KCC2*^*E/+*^ mice spent more time to explore the novel stranger compartment than the familiar stranger compartment (**Fig. S3A**, *P*=0.007, n=14 for *KCC2*^*+/+*^ and *P*=0.008, n=14 for *KCC2*^*E/+*^ mice, one-way Anova test), and there were no difference between *KCC2*^*+/+*^ and *KCC2*^*E/+*^ mice in the number of entries in the 3 different chambers compartment (**Fig. S3B**, *P*=0.3, n=76,70, two-way Anova test). Two other behavioral tests that are used to determine anxiety behavior, locomotion, and depression revealed no difference in the behavior of *KCC2*^*+/+*^ and *KCC2*^*E/+*^ mice (**Fig. S4, Table S3**). These data show that KCC2 T906E/T1007E impaired communication and sociability behavior.

### Bumetanide restores altered glutamate/GABA balance and seizure susceptibility, but not vocalization and social interactions, in *KCC2*^*E/+*^ mice

Prenatal (Tyzio *et al.*, 2014) or adult (Deidda *et al.*, 2015; Dargaei *et al.*, 2018) administration of the NKCC1 inhibitor bumetanide has been shown to alleviate symptoms in rat and mice animal models of NDDs, presumably due to rescuing of depolarizing action of GABA (Deidda *et al.*, 2014). We therefore examined whether bumetanide could restore neuronal network dysfunction and correct behavioral alterations in *KCC2*^*E/+*^ mice.

To this aim, *KCC2*^*E/+*^ mice received daily sub-cutaneous injection (0.2mg/kg) from P6 to P15, a period during which the hyperpolarizing shift of GABA responses occurs in *KCC2*^*+/+*^ mice but failed in *KCC2*^*E/+*^ mice. We found that the bumetanide treatment fully restored the E/I balance (**Fig. 6A**). The sGlut-PSCs to sGABA-PSCs ratio was similar between the naïve and bumetanide-treated *KCC2*^*+/+*^ mice (0.5±0.05, n=11 neurons from 3 mice [n=11,3], *P*=0.3, one-way Anova test) but significantly lower in bumetanide-treated *KCC2*^*E/+*^ mice compared to sham *KCC2*^*E/+*^ mice (0.6±0.1 [n=13,3] vs 2.6±0.8 [n=12,3], *P*=0.01, one-way Anova test). Next, to determine whether the rescue effect of bumetanide relies on the restoration of the GABA hyperpolarizing shift or on the reorganization of neuronal circuits, we assessed the effect of acute bumetanide treatment *in vitro*. 3h treatment of P15 hippocampal slices with bumetanide (10µM) fully restored the sGlut-PSCs to sGABA-PSCs ratio of CA3 pyramidal neurons (**Fig. 6A**, from 2.6±0.8 in control *KCC2*^*E/+*^ slice [n=12, 3] vs 0.7±0.1 in bumetanide-treated *KCC2*^*E/+*^ slice [n=10, 3], *P*=0.04, one-way Anova test).

**Figure 6.**
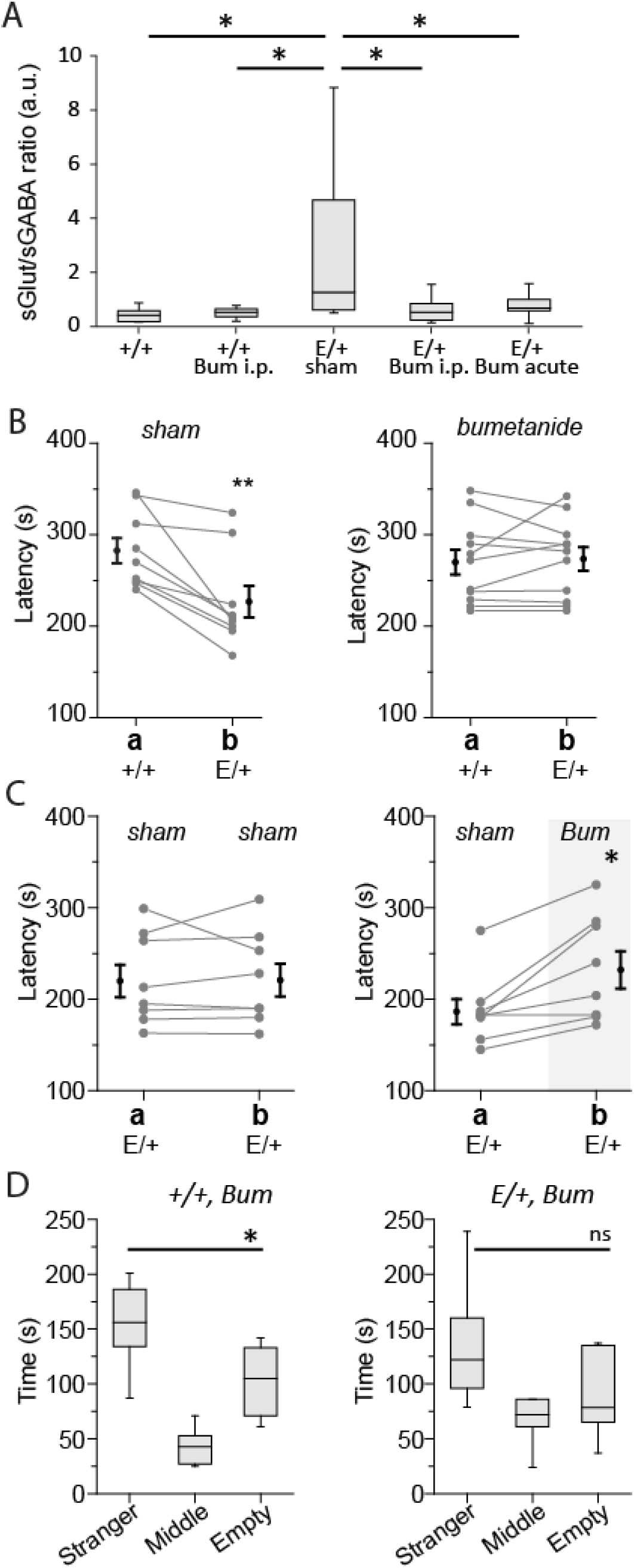
Bumetanide treatment restores the Glutamate/GABA balance in *KCC2E/+* mice. Bumetanide (bum) or DMSO (sham) was administrated i.p. daily from P6 to P15 or applied by bath (acute). **A)** Box plot of the frequency ratio of spontaneous glutamatergic to GABAergic postsynaptic currents. *P<0.05, one-way Anova test. Numbers in parenthesis indicate the number of cells recorded and mice used. **B)** Latency of the onset of tonic-clonic seizures of *KCC2*^*+/+*^ and *KCC2*^*E/+*^ mice in compartments **a** and **b** and treated with DMSO (vehicle, left plot) or bumetanide (right plot). The connected points indicate pairs of animals. The points with error bars indicate mean ± SEM values. **P<0.01, Wilcoxon paired test. **C)** *KCC2*^*E/+*^ mice treated with bumetanide (bum, compartment **b**, right plot) exhibit longer latencies of seizures onset compared to DMSO treated mice (compartment **a**, left plot). Left plot shows control experiments when **a** and **b** compartments contained *KCC2*^*E/+*^ mice treated with DMSO (veh). The connected points indicate pairs of animals. The points with error bars indicate mean ± SEM values. *P<0.05, Wilcoxon Paired test. **D)** Social behavior of *KCC2*^*+/+*^ (n=10) and *KCC2*^*E/+*^ (n=11) mice treated with bumetanide from P6 to P15 and studied at P60. ns= non-significant, *P<0.05, one-way Anova test).

Chronic bumetanide administration from P6 to P15 in P15 *KCC2*^*E/+*^ mice fully restored the latency of flurothyl-induced tonic-clonic seizures as compared with *KCC2*^*+/+*^ littermates (**Fig. 6B**, right plot, *P*=0.6, 11 pairs). The observed effect was specific for bumetanide, as *KCC2*^*E/+*^ animals treated from P6 to P15 with vehicle only (DMSO) showed similar shorter latencies of seizures induction (**Fig. 6B**, left plot, *P*=0.02, 9 pairs). Consistent with above observation, P15 *KCC2*^*E/+*^ mice treated with bumetanide showed significantly longer latencies of seizure responses as compared to their *KCC2*^*E/+*^ littermates treated with vehicle (**Fig. 6C**, right plot, *P*=0.016, 8 pairs), while vehicle-treated *KCC2*^*E/+*^ mice placed into two compartments showed similar seizure latencies (**Fig. 6C**, left plot, *P*=0.61, 8 pairs).

In contrast, chronic bumetanide treatment from P6 to P15 did not normalize the social behavior deficits observed in *KCC2*^*E/+*^ mice tested at P60. *KCC2*^*E/+*^ mice treated with bumetanide spent as much time exploring the empty compartment and the compartment with strangers (**Fig. 6D**, *P*= 0.13, n=7, one-way Anova test) as did untreated *KCC2*^*E/+*^ mice (**Fig. 5D**). As control experiments, *KCC2*^*+/+*^ mice were treated with bumetanide from P6 to P15. Like *KCC2*^*+/+*^*mice*, bumetanide-treated *KCC2*^*+/+*^ mice spent more time to explore stranger compartment than empty compartment (**Fig. 6D**, *P*=0.003, n=10, one-way Anova test).

## DISCUSSION

By employing a multidisciplinary approach that includes biochemistry, electrophysiology, and neurobehavior in Thr^906^/Thr^1007^ phospho-mimetic *KCC2*^*E/+*^ mice, we have made several novel observations that illustrate the importance of KCC2 Thr^906^/Thr^1007^ phosphorylation for post-natal brain physiology and neurodevelopment. We have shown that P15, but not P30, *KCC2*^*E/+*^ mice exhibit an alteration in strength of GABAergic inhibition and an enhanced excitatory/inhibitory (E/I) ratio, two characteristic electrophysiological signatures associated with multiple NDDs (e.g., (Rubenstein and Merzenich, 2003; Cellot and Cherubini, 2014; Deidda *et al.*, 2014)). Furthermore, behavioral tests show phospho-mimetic KCC2 Thr^906^/Thr^1007^ mutation lead to increased ultra-sonic vocalization during the two first post-natal weeks, higher susceptibility for seizure generation at P15, and impaired social interaction in P60 mice, three characteristic neurobehaviors associated with multiple NDDs (e.g., (Scattoni *et al.*, 2009; Bey and Jiang, 2014; Pasciuto *et al.*, 2015)). Post-natal treatment with the NKCC1 blocker bumetanide restored E/I balance and normalized seizure thresholds but failed to restore the social interaction impairments. These data highlight the functional importance of KCC2’s post-translational control in CNS development, and suggest impairment of this mechanism may contribute to the pathogenesis of certain symptoms associated with multiple different NDDs.

### Delayed emergence of GABAergic inhibition in *KCC2*^*E/+*^ mice

By performing electrophysiological recordings in acute hippocampal slices, we showed that the developmental maturation of Cl^-^-dependent GABAergic neurotransmission is delayed in *KCC2*^*E/+*^ mice. At P12-P15, DF_GABA_ is shifted towards negative values in *KCC2*^*+/+*^ mice, but remained positive in *KCC2*^*E/+*^ mice. Likewise, at P12, isoguvacine reduced the firing of CA3 pyramidal cell in *KCC2*^*+/+*^ mice, but enhanced the firing in *KCC2*^*E/+*^ mice. These results corroborate previous *in vitro* studies showing that phospho-mimetic mutations of KCC2 Thr^906^/Thr^1007^ down-regulate transporter activity, leading to an increase in [Cl^-^]_i_, and consequently, reduced inhibitory strength of GABA (Friedel *et al.*, 2015). Interestingly, the developmental action of GABA in *KCC2*^*E/+*^ mice was not abolished but delayed; at P30, the DF_GABA_ and effect of isoguvacine were similar in *KCC2*^*+/+*^ and *KCC2*^*E/+*^ animals. This could be explained by compensation of the KCC2 T906E/T1007E allele by the developmental up-regulation in KCC2 abundance and simultaneous Thr^906^/Thr^1007^ de-phosphorylation in the wild type KCC2 allele. Together, these results indicate that post-translational modification of KCC2 is a determining factor in the post-natal emergence of functional GABAergic inhibition.

### KCC2 post-translational control of network function and seizure susceptibility

In addition to causing a reduction in the inhibitory strength of GABA, the T906E/T1007E phospho-mimetic mutations of KCC2 caused a transient enhancement of network excitability and the susceptibility to generate seizures. All three parameters (DF_GABA_, the E/I ratio, and seizure susceptibility) are altered at P15, but not P30, indicating a correlation between the polarity of GABA, an increase in neuronal network activity, and the sensitivity to epileptogenic agents. Furthermore, chronic post-natal treatment with bumetanide rescued the E/I ratio and alleviated the sensitivity to epileptogenic agents. Interestingly, the same rescue of E/I ratio was observed after acute bumetanide application to hippocampal slices. Thus, the increase in network excitability and seizure susceptibility in *KCC2*^*E/+*^ mice likely results from delayed depolarizing action of GABA and not from abnormal developmental sequences or reorganization of neuronal networks. Our findings also corroborate and extend recent work showing that genetic mutation of Thr 906/1007 to alanine (Ala), the *inverse* of our phospho-mimetic model that mimics constitutive dephosphorylation, enhances KCC2 activity and limits the onset and severity of seizures in homozygous mice (Moore *et al.*, 2018).

### KCC2 post-translational control and NDDs

We have shown that *KCC2*^*E/+*^ mice exhibit behavioral alterations, including an increase in the number of ultrasonic calls emitted by P10 and P12 isolated pups, and reduced social interactions in P60 mice, two key symptoms of ASDs. There was no difference in tests that evaluated anxiety, locomotion, and depression, although detailed investigations are needed to confirm these results. Previous work has implicated intrinsic KCC2 malfunction or dysregulation in different epilepsy subtypes (Palma *et al.*, 2006; Huberfeld *et al.*, 2007; Talos *et al.*, 2012; Kahle *et al.*, 2014; Puskarjov *et al.*, 2014), and multiple NDDs such as Schizophrenia, ASDs, Rett syndrome (Arion and Lewis, 2011; Tyzio *et al.*, 2014; Merner *et al.*, 2015; Sullivan *et al.*, 2015; Banerjee *et al.*, 2016; Tang *et al.*, 2016); The present work is the first study demonstrating that the post-translational KCC2 control could be a risk factor in the NDDs etiology. Our findings extend several recent genetic studies that have identified mutation-linked modifications of the post-translational control of KCC2 in human neurological disorders (Kahle *et al.*, 2014; Puskarjov *et al.*, 2014; Stödberg *et al.*, 2015). Interestingly, post-natal treatment with bumetanide in our mice model failed to rescue interaction social behavioral deficit. Thus, adult social interaction alteration does not depend on the post-natal developmental GABAergic sequence.

### Link between electrophysiological alteration and neuro-behavioral modification

Impaired GABAergic neurotransmission and network activity dysregulation are hallmarks of NDD (Rubenstein and Merzenich, 2003; Cellot and Cherubini, 2014; Deidda *et al.*, 2014). Impaired Cl-homeostasis and excitatory GABA activity have been reported in adult mice model of Rett syndrome (Banerjee *et al.*, 2016), Down syndrome (Deidda *et al.*, 2015) and Huntington’s disease (Dargaei *et al.*, 2018). In these models, bumetanide treatment in adults restored the hyperpolarizing action of GABA and alleviated the symptoms. In contrast, in our study, the alterations of social interaction are present while GABA has shifted toward hyperpolarizing direction. Thus, the *KCC2*^*E/+*^ mice is a valuable model to study the patho-physiological importance of post-natal Cl-modification and neuronal network activity.

Previous studies have suggested dysregulation of the post-natal GABAergic sequence contributes to the pathogenesis of several neurological disorders by impairing neuronal network formation (Tyzio *et al.*, 2006, 2014; Eftekhari *et al.*, 2014; He *et al.*, 2014). Consistent with this, hippocampal neurons of P0-P30 age Fragile X and Valproate mouse models of ASD exhibit increased post-natal Cl^-^ and neuronal network activity (Eftekhari *et al.*, 2014; He *et al.*, 2014; Tyzio *et al.*, 2014). In these models, peri-natal treatment with bumetanide restored the GABAergic post-natal sequence (Tyzio *et al.*, 2014) and adult behavior alterations (Eftekhari *et al.*, 2014). In our experiments, the P6-P15 post-natal treatment of *KCC2* ^*E/+*^ animals with bumetanide restored neuronal network activity at P15-P20, but failed to rescue compromised social behavior at P60. Taken together, these data indicate that the post-natal alteration of GABAergic transmission and network functioning is not implicated in social behavior modification and alteration of this phenomenon in *KCC2* ^*E/+*^ mice involves other KCC2-dependent mechanism. Contrary to *KCC2*^*E/+*^ monogenic model, all cited above works were performed on multifactorial models involving long-lasting changes of large number of genes and signaling pathways (Bey and Jiang, 2014; De Rubeis *et al.*, 2014; Ehrhart *et al.*, 2016) as well as reported changes of neuronal Cl^-^ homeostasis in brain slices from juvenile (P30, (Tyzio *et al.*, 2014)) or young adult (P60, (Deidda *et al.*, 2015; Dargaei *et al.*, 2018)) animals.

### Clinical relevance of *KCC2*^*E/+*^ mice model

Recent clinical studies revealed that heterozygous mutations in human KCC2 are associated with multiple neurological disorders featuring impaired Cl^-^ homeostasis including several forms of epilepsy (Kahle *et al.*, 2014; Puskarjov *et al.*, 2014; Stödberg *et al.*, 2015; Saitsu *et al.*, 2016), ASD, and schizophrenia (Merner *et al.*, 2015). How these mutations affect KCC2 activity and/or expression in humans and whether these mutations act in combination with other risk factors is presently unknown. *In vitro* experiments performed on cell cultures expressing KCC2 harboring human mutations suggest that at least some of these mutations impair critical post-translational modifications of KCC2 that alter its activity (Kahle *et al.*, 2014; Stödberg *et al.*, 2015). Our study supports these genetic findings by showing that impaired regulation of KCC2 phosphorylation in even one allele can directly contribute to the formation of pathology relevant for multiple NDDs. These KCC2 T906E(A)/T1007E(A) transgenic animals thus represent valuable models to dissect the importance and effects of regulated KCC2 phosphorylation *in vivo*, and serve as genetic proof that drug development targeting these sites is a compelling strategy to powerfully modulate KCC2 activity.

## Author contribution

L.I.P., J-L.G. and I.M. designed research, performed experiments and wrote the paper. K.T.K. designed research and wrote the paper. J.Z., D.D. and I.K. performed experiments. J.D., L.I.P., I.M. bred the colony.

## ACKNOWLEDGMENTS

We express our gratitude to the Neurochlore team (www.neurochlore.fr) for their help in the mouse neurobehaviour studies; Mrs Marie Kurz for help in animal care; and Mrs Aurélie Montheil and Mrs Francesca Bader for genotyping.

