## Supplementary for "Impaired KCC2 phosphorylation leads to neuronal network dysfunction and neurodevelopmental pathogenesis"

### SUPPLEMENTARY MATERIAL

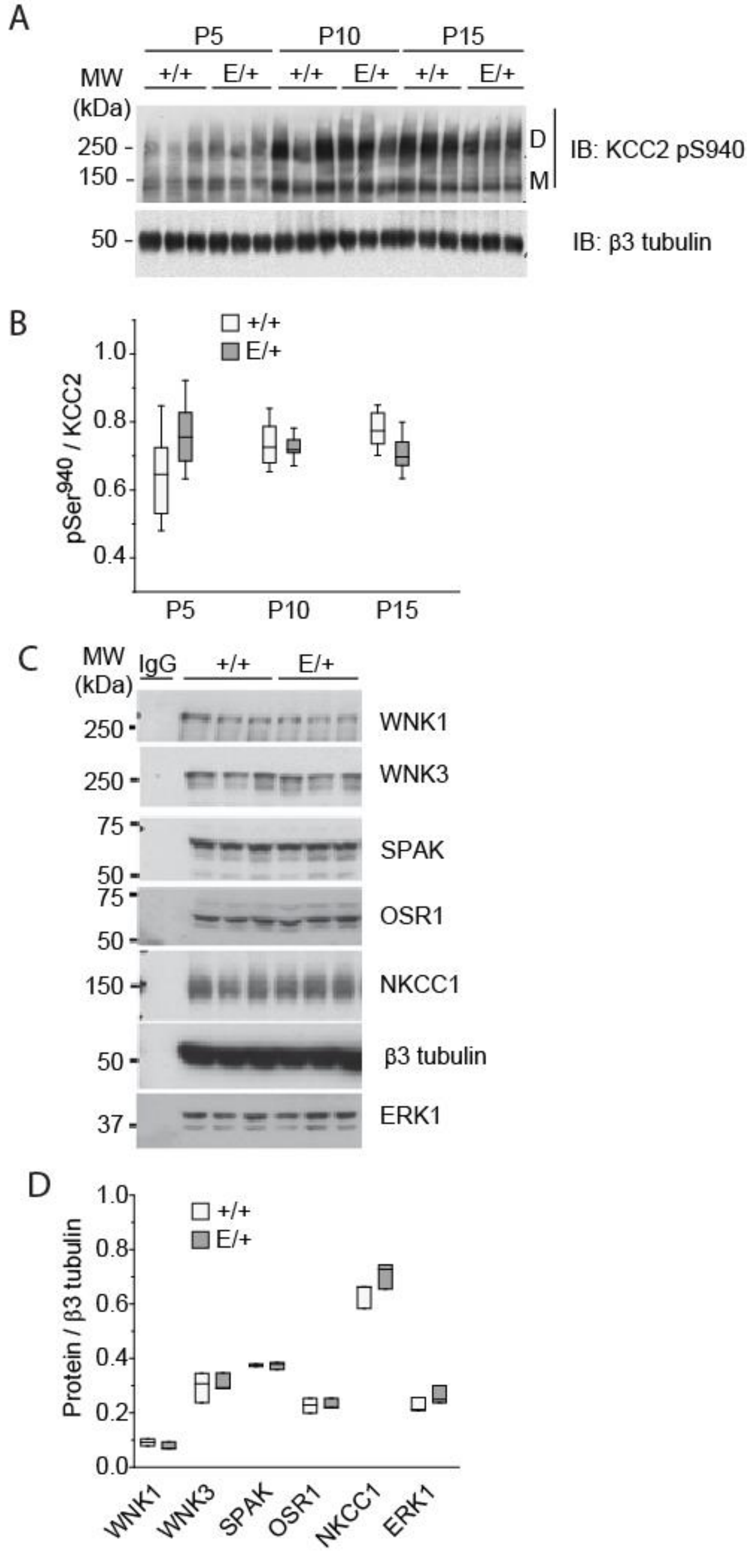


**Figure S1: KCC2 related signaling in KCC2$^{E/+}$ phospho-mimetic mice.**

**A)** Developmental expression of phosphorylated Ser$^{940}$ pool (pS940) of KCC2 in *KCC2$^{+/+}$* and *KCC2$^{E/+}$* pups. Hippocampi lysates from in *KCC2$^{E/+}$* and *KCC2$^{+/+}$* mice at the indicated developmental points were subjected for Western blots and revealed using anti-pSer940 antibody. An antibody recognizing neuron specific β3 tubulin was employed to normalize samples loading. The same set of samples as shown in Fig.1 D. Each condition illustrates migration of extracts from 3 mice. **B)** Boxplot and individual values of the abundance of pS940 normalized to the total expression of KCC2 illustrated in Fig.1 D. The two-way Anova test revealed neither statistically significant difference between *KCC2$^{E/+}$* and *KCC2$^{+/+}$* mice nor during development. **C)** Western blots of P15 hippocampi lysates from three *KCC2$^{E/+}$* and three *KCC2$^{+/+}$* mice revealed using different KCC2 metabolism – related antibodies. **D)** Statistical boxplots of quantified abundancies of indicated proteins. Non-parametric Man-Whitney test revealed no difference between *KCC2$^{E/+}$* and *KCC2$^{+/+}$* mice for all studied proteins. (*P*=0.38, n=3 for WNK1; *P*=0.99, n=3 for WNK3; *P*=0.82, n=3 for SPAK; *P*=0.98, n=3 for OSR1; *P*=0.38, n=3 for NKCC1; *P*=0.38, n=3 for ERK1.

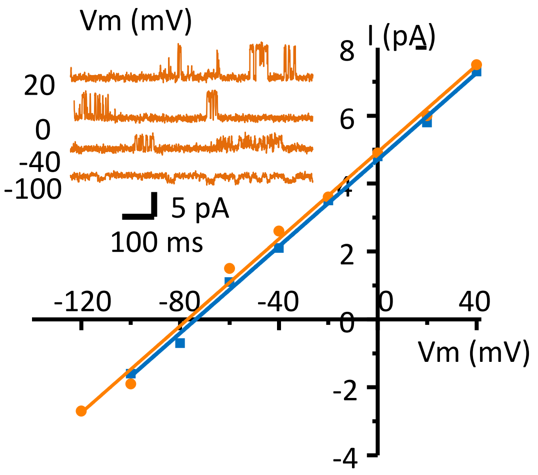


**Figure S2: Similar resting membrane potential in P13 CA3 neurons of acute slices from *KCC2*$^{E/+}$ and *KCC2*$^{+/+}$ mice.** Traces illustrate representative single NMDA receptor-channels currents recorded at different potentials from CA3 neurons of *KCC2*$^{E/+}$ mice. Representative I-V curves of single NMDA receptor-channels currents recorded from CA3 neurons of *KCC2*$^{E/+}$ (brown) and *KCC2*$^{+/+}$ (blue) mice.

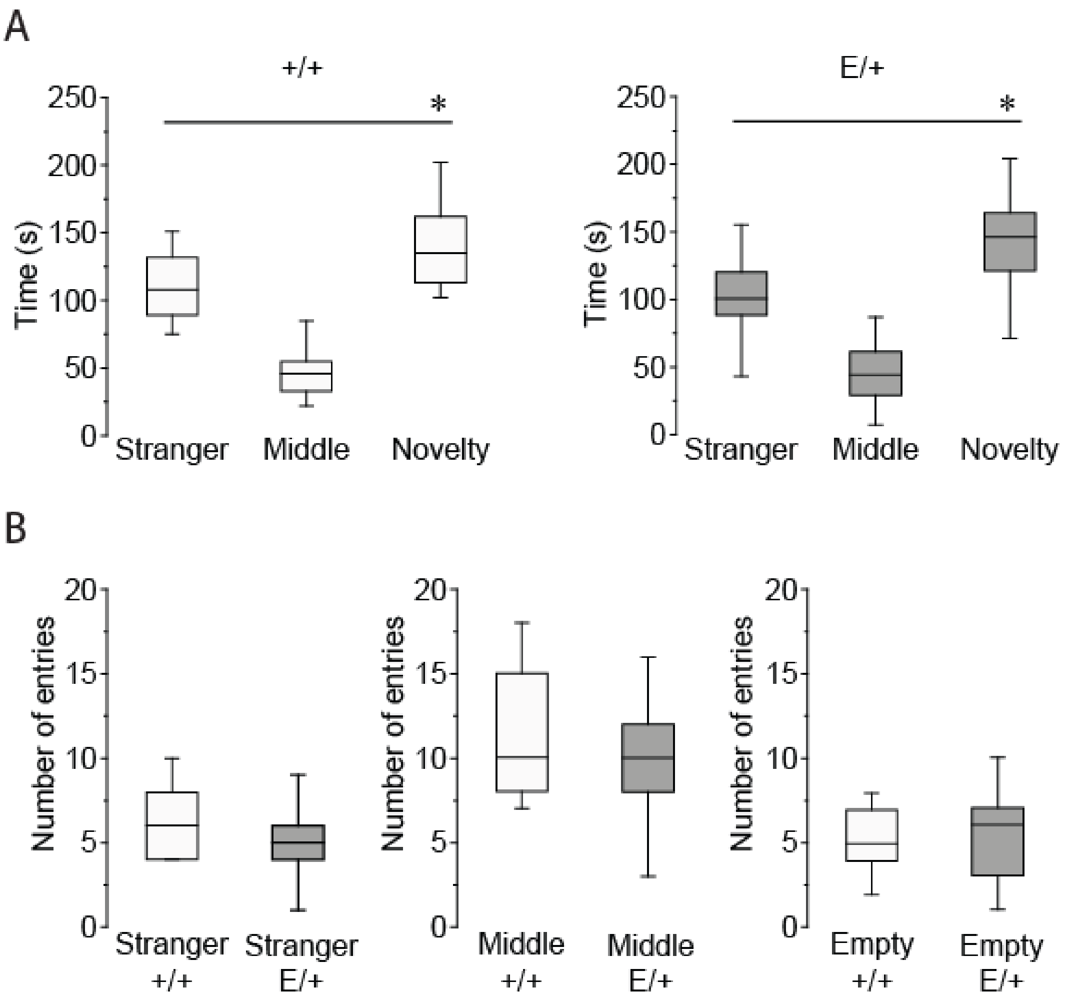


**Figure S3: Mutations T906E/T1007E do not modify social novelty (A) and exploration (B) behavior.**

**A)** Time spent by *KCC2*$^{+/+}$ (left plots) and *KCC2*$^{E/+}$ (right plot) mice in stranger and novelty chambers. ns= non-significant, *$P$<0.05, two-tailed unpaired Student's t-test. **B)** There was no statistically significant difference in number of entries to each chamber of *KCC2*$^{+/+}$ and *KCC2*$^{E/+}$ mice. P>0.05 for all cases, two-tailed unpaired Student's t-test.

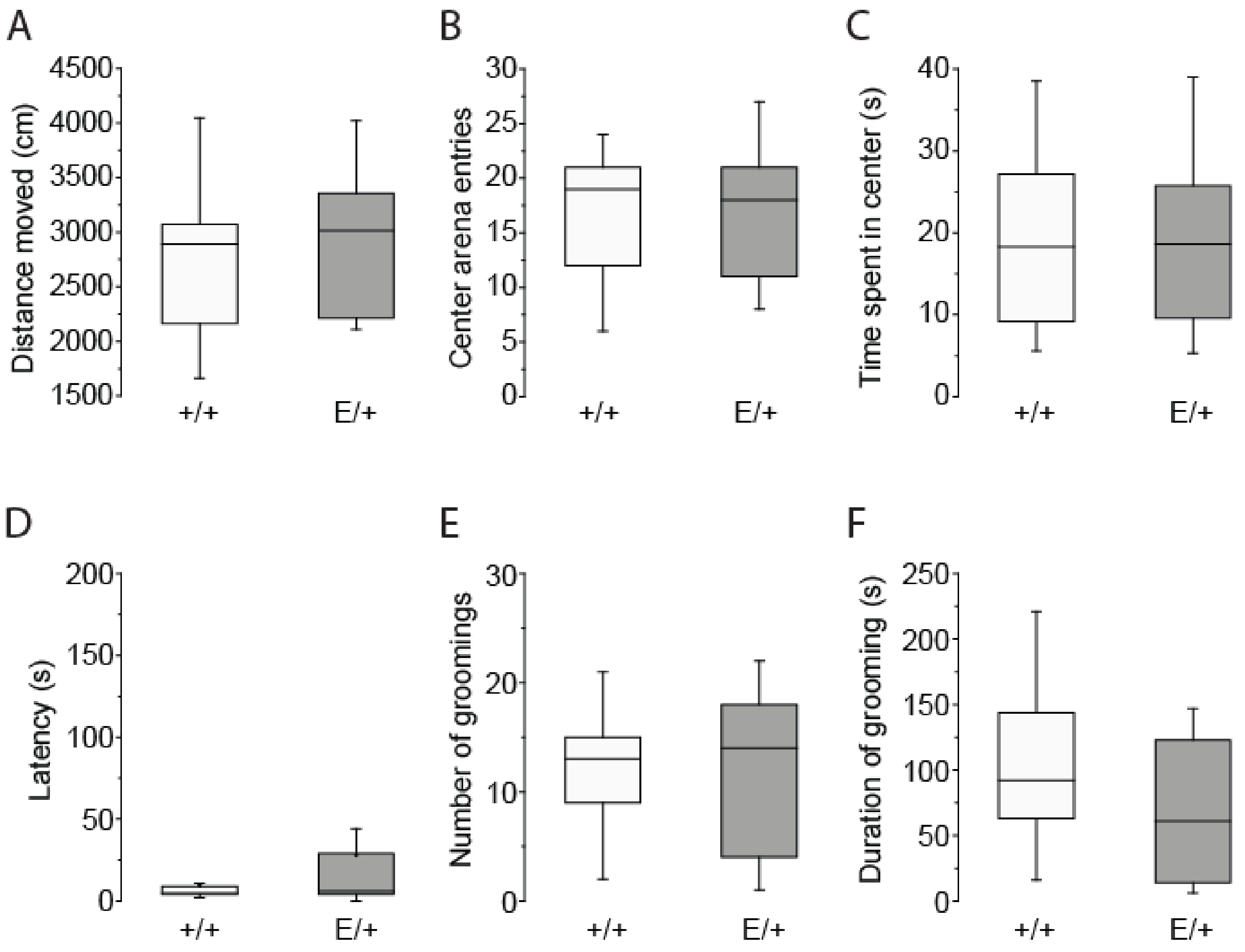


**Figure S4: Results of additional behavior tests in KCC2$^{+/+}$ (WT) and KCC2$^{E/+}$ (E/+) mice.**
**A)** Locomotion test assayed using open field. Distances run by mice in arena. **B,C)** Anxiety test in open field. Number of entries (B) and time spent (C) by mice in the central zone. **D,E,F)** Depressive like behavior of KCC2$^{E/+}$ and KCC2$^{+/+}$ assayed using splash test. Latency to start grooming (D), number of grooming events (E), and the duration of grooming (F). None of tests showed statistically significant difference between behavior of *KCC2*$^{+/+}$ and *KCC2*$^{E/+}$ mice. $P>0.05$, two-tailed unpaired Student's t-test.

**Supplementary Table S1: Statistical differences among the samples illustrated in Figures 1 and S1.**

| Data reference | Data structure | Type of test | Power |
|---|---|---|---|
| Fig.1 B +/+ vs E/+ | Normal distribution | Two-way Anova | $F$ (1,279)=1.84, $P$ =0.18 |
| Fig.1 B age dependence | Normal distribution | Two-way Anova | $F$ (9,279)=205.39, $P$ =1.1E-8 |
| Fig.1 C +/+ vs E/+ | Non-normal distribution | Chi-square test | $X^2$ (2, n=396)=4.97, $P$ =0.83 |
| Fig.1 E, +/+ vs E/+ | Normal distribution | Two-way Anova | $F$ (1,32)=29.6, $P$ =5.5E-6 |
| Fig.1 E, age dependence | Normal distribution | Two-way Anova | $F$ (2,32)=24.0, $P$ =4.3E-7 |
| P5, +/+ vs E/+ | | post-hoc Tukey | $P$ =7.5E-5 |
| P10, +/+ vs E/+ | | post-hoc Tukey | $P$ =0.008 |
| P15, +/+ vs E/+ | | post-hoc Tukey | $P$ =0.09 |
| Fig.1 F, +/+ vs E/+ | Normal distribution | Two-way Anova | $F$ (1,32)=29.6, $P$ =5.4E-6 |
| Fig.1 F, age dependence | Normal distribution | Two-way Anova | $F$ (2,107)=24.0, $P$ =4.3E-7 |
| P5, +/+ vs E/+ | | post-hoc Tukey | $P$ =0.002 |
| P10, +/+ vs E/+ | | post-hoc Tukey | $P$ =0.001 |
| P15, +/+ vs E/+ | | post-hoc Tukey | $P$ =0.03 |
| Fig.1 G, +/+ vs E/+ | Normal distribution | Two-way Anova | $F$ (1,32)=1.33, $P$ =0.26 |
| Fig.1 G, age dependence | Normal distribution | Two-way Anova | $F$ (2,32)=94.5, $P$ =3.7E-14 |
| P5, +/+ vs E/+ | | post-hoc Tukey | $P$ =0.81 |
| P10, +/+ vs E/+ | | post-hoc Tukey | $P$ =0.37 |
| P15, +/+ vs E/+ | | post-hoc Tukey | $P$ =0.73 |
| Fig.S1B | Non-normal distribution | Mann-Whitney U-test | |
| WNK1+/+ vs WNK1 E/+ | | | $U$ =7, n=3,3; $p$ =0.38 |
| WNK3 +/+ vs WNK3 E/+ | | | $U$ =4, n=3,3; $p$ =0.98 |
| SPAK +/+ vs SPAK E/+ | | | $U$ =3.5, n=3,3; $p$ =0.82 |
| OSR1 +/+ vs OSR1 E/+ | | | $U$ =4.5, n=3,3; $p$ =0.99 |
| NKCC1 +/+ vs NKCC1 E/+ | | | $U$ =2, n=3,3; $p$ =0.38 |
| ERK1 +/+ vs ERK1 E/+ | | | $U$ =2, n=3,3; $p$ =0.38 |
| Fig. S1D, +/+ vs E/+ | Normal distribution | Two-way Anova | $F$ (1,32)=0.16, $P$ =0.69 |
| Fig. S1D, age dependence | Normal distribution | Two-way Anova | $F$ (2,32)=0.53, $P$ =0.59 |
| P5, +/+ vs E/+ | | post-hoc Tukey | $P$ =0.12 |
| P10, +/+ vs E/+ | | post-hoc Tukey | $P$ =0.74 |
| P15, +/+ vs E/+ | | post-hoc Tukey | $P$ =0.06 |

**Supplementary Table S2: Statistical differences among the samples illustrated in Figure 2.**

| Data reference | Data structure | Type of test | Power |
|---|---|---|---|
| Fig.2 B, +/+ | Normal distribution | One-way Anova | *F* (3,71)=14.1, *P* =2.6E-07 |
| P7-9 vs P12-14 | | post-hoc Tukey | *P* =1.26E-5 |
| P7-9 vs P17-20 | | post-hoc Tukey | *P* =9.46E-6 |
| P12-14 vs P17-20 | | post-hoc Tukey | *P* =0.98 |
| P17-20 vs P30 | | post-hoc Tukey | *P* =0.93 |
| Fig.2 B, E/E | Normal distribution | One-way Anova | *F* (3,71)=25.9, *P* =2.0E-11 |
| P7-9 vs P12-14 | | post-hoc Tukey | *P* =0.87 |
| P7-9 vs P17-20 | | post-hoc Tukey | *P* =4.0E-4 |
| P12-14 vs P17-20 | | post-hoc Tukey | *P* =0.003 |
| P17-20 vs P30 | | post-hoc Tukey | *P* =0.03 |
| Fig.2 B +/+ vs E/+ | Normal distribution | Two-way Anova | *F* (1,145)=17.2, *P* =5.60E-5 |
| Fig.2 B age dependence | Normal distribution | Two-way Anova | *F* (3,145)=25.3, *P* =5.18E-13 |
| P7-9, +/+ vs E/+ | | post-hoc Tukey | *P* =0.33 |
| P12-14, +/+ vs E/+ | | post-hoc Tukey | *P* =5.7E-7 |
| P17-20, +/+ vs E/+ | | post-hoc Tukey | *P* =0.002 |
| P30, +/+ vs E/+ | | post-hoc Tukey | *P* =0.5 |

| | | | |
|---|---|---|---|
| Fig.2 C +/+ vs E/+ | Normal distribution | Two-way Anova | *F* (1,33)=0.53, *P* =0.47 |
| Fig.2 C age dependence | Normal distribution | Two-way Anova | *F* (3,33)=1.92, *P* =0.15 |

| | | | |
|---|---|---|---|
| Fig.2 E +/+ vs E/+ | Non-normal distribution | Chi-square test | $X^2$ (2, n=76)=44.3, *P* =1E-5 |

| | | | |
|---|---|---|---|
| Fig.2 F, +/+ vs E/+ | Normal distribution | Two-way Anova | *F* (1,107)=19.8, *P* =2.16E-5 |
| Fig.2 F, age dependence | Normal distribution | Two-way Anova | *F* (3,107)=18.0, *P* =1.54E-9 |
| P6-9, +/+ vs E/+ | | post-hoc Tukey | *P* =0.99 |
| P10-13, +/+ vs E/+ | | post-hoc Tukey | *P* =0.045 |
| P14-17, +/+ vs E/+ | | post-hoc Tukey | *P* =0.009 |
| P18-21, +/+ vs E/+ | | post-hoc Tukey | *P* =0.046 |
| P30, +/+ vs E/+ | | post-hoc Tukey | *P* =0.08 |

**Supplementary Table S3: Statistical differences among the samples in behavior experiments illustrated in Figures 5, S3 and S4.**

| Data reference | Data structure | Type of test | Power |
|---|---|---|---|
| Fig.5 A, +/+ vs E/+ | Normal distribution | Two-way Anova | *F*(1,180)=9.08, *P*=0.003 |
| Fig.5 A age dependence | Normal distribution | Two-way Anova | *F*(4,180)=8.73, *P*=1.83E-6 |
| P2, +/+ vs E/+ | | post-hoc Tukey | *P=0.19* |
| P4, +/+ vs E/+ | | post-hoc Tukey | *P=0.86* |
| P8, +/+ vs E/+ | | post-hoc Tukey | *P=0.73* |
| P10, +/+ vs E/+ | | post-hoc Tukey | *P=0.005* |
| P12, +/+ vs E/+ | | post-hoc Tukey | *P=0.003* |
| Fig.5 C, +/+ | Normal distribution | One-way Anova | |
| Stranger vs Empty | | | *F(1,28)=16.4, P=3.68E-4* |
| Stranger vs Middle | | | *F(1,28)=72.97, P=2.76E-9* |
| Empty vs Middle | | | *F(1,28)=25.56, P=2.38E-5* |
| Fig.5 D, E/+ | Normal distribution | One-way Anova | |
| Stranger vs Empty | | | *F(1,26)=2.85, P=0.10* |
| Stranger vs Middle | | | *F(1,26)=50.33, P=1.56E-7* |
| Empty vs Middle | | | *F(1,26)=18.16, P=2.36E-4* |
| Fig.S3A, +/+ | Normal distribution | One-way Anova | |
| Stranger vs Empty | | | *F(1,26)=8.66, P=0.007* |
| Stranger vs Middle | | | *F(1,26)=51.10, P=1.36E-7* |
| Empty vs Middle | | | *F(1,26)=90.13, P=6.18E-10* |
| Fig.S3A, E/+ | Normal distribution | One-way Anova | |
| Stranger vs Empty | | | *F(1,26)=8.27, P=0.008* |
| Stranger vs Middle | | | *F(1,26)=21.47, P=8.85E-5* |
| Empty vs Middle | | | *F(1,26)=64.75, P=1.59E-8* |
| Fig.S3B, +/+ | Normal distribution | One-way Anova | |
| Stranger, +/+ vs E/+ | | | *F(1,27)=2.45, P=0.13* |
| Middle, +/+ vs E/+ | | | *F(1,27)=0.38, P=0.56* |
| Empty, +/+ vs E/+ | | | *F(1,27)=0.06, P=0.81* |
| Fig.S4 A, B, C | Normal distribution | One way Anova-test | |
| Number of grooming | | | *F(1,30)=0.002, P=0.96* |
| Duration of grooming | | | *F(1,30)=2.7, P=0.11* |
| Latency | | | *F(1,30)=2.1, P=0.15* |
| Fig.S4 D, E, F | Normal distribution | One way Anova-test | |
| Distance moved | | | *F(1,30)=0.7, P=0.42* |
| Center area entries | | | *F(1,30)=0.04, P=0.83* |
| Time spent in center | | | *F(1,30)=0.002, P=0.96* |

**Supplementary Table S4: Statistical differences among the samples illustrated in Figure 6 D.**

| Data reference | Data structure | Type of test | Power |
|---|---|---|---|
| Fig.6 D, +/+ Bum | Normal distribution | One-way Anova | |
| Stranger vs Empty | | | *F(1,18)=11.78, P=0.003* |
| Stranger vs Middle | | | *F(1,18)=76.82, P=6.52E-8* |
| Empty vs Middle | | | *F(1,18)=27.90, P=5.07E-5* |
| Fig.6 D, E/+ Bum | Normal distribution | One-way Anova | |
| Stranger vs Empty | | | *F(1,10)=2.70, P=0.13* |
| Stranger vs Middle | | | *F(1,10)=4.64, P=0.06* |
| Empty vs Middle | | | *F(1,10)=0.36, P=0.56* |
